# Validated CRISPR/Cas9 guide RNAs targeting neurodevelopmental genes in the tunicate *Ciona robusta*

**DOI:** 10.64898/2026.03.25.711585

**Authors:** Sydney Popsuj, Tenzin Kalsang, Kwantae Kim, Erica Drummond, Pooja Manekar, Pranavvarma Munagapati, Manasi Oleti, Hiroki Sato, Izabella Vickery, Eduardo D. Gigante, Alberto Stolfi

## Abstract

The development of the central nervous system (CNS) depends on tightly regulated gene expression programs that guide neural progenitor differentiation and neuronal subtype specification. The tunicate *Ciona robusta* provides a powerful and simplified model for dissecting the genetic control of nervous system development, with a larval CNS composed of just over 200 neurons and sensory cells. Although CRISPR/Cas9-mediated mutagenesis is now routinely used in *Ciona*, validated single-guide RNAs (sgRNAs) have yet to be validated for key neural genes. Here, we report the design and experimental validation of 25 novel sgRNAs targeting eight conserved genes encoding conserved proteins involved in neurodevelopment and neural function, including six transcription factors (*Cdx, Foxb, Sox1/2/3, Dmbx, Engrailed*, and *Mnx*) and two neural effector genes (*Tyrosinase* and *Slc18a3/VAChT*). Candidate sgRNAs were selected using CRISPOR and tested for mutagenesis efficiency using Illumina-based target site amplicon sequencing. All sgRNAs induced insertions or deletions at their target loci, with most genes yielding at least one sgRNA with mutagenesis efficacy exceeding 30%, with the exception of *Dmbx*, for which maximal efficacy reached 25%. We further compared measured mutagenesis rates with predicted Doench ‘16 and Doench Ruleset 3 (RS3) scores, observing a modest but improved correlation with RS3 predictions. Based on these results, we recommend considering both scoring algorithms, with RS3 potentially offering improved predictive value for *Ciona*.

## Introduction

The development of the central nervous system (CNS) in any organism relies on the precise spatial and temporal regulation of genes that regulate the differentiation of neural progenitors into distinct, fully functional neuronal subtypes. Transcription factors contribute to diverse functions in development of the CNS, such as cell cycle regulation, signal transduction, and establishing proper neuronal connectivity, through their transcriptional regulation of target genes encoding downstream regulators and effectors of these processes (Hobert and Kratsios, 2019). *Ciona robusta*, a tunicate, represents a simplified model system for investigating nervous system development, with a much reduced CNS comprised of little over 200 neurons and sensory cells (Nishino, 2018; Ryan and Meinertzhagen, 2019).

Like many tunicates, *Ciona* follow a biphasic lifecycle, metamorphosing from a swimming larva into a sessile adult (Karaiskou et al., 2015). Swimming behaviors in *Ciona* are directly controlled by a handful of CNS cells that comprise the motor ganglion (MG) and tail nerve cord, parts of which might be homologous to vertebrate hindbrain and spinal cord (Kourakis et al., 2025; Piekarz and Stolfi, 2024). Various swimming behaviors enable the larvae to identify suitable, hard substrates on which to settle and initiate metamorphosis. These include different chemotactic, rheotactic, phototactic and geotactic behaviors that depend on relatively simple circuits connected to sensory cells in the epidermis or the larval brain vesicle of the CNS (Bostwick et al., 2019; Hoyer et al., 2024; Kourakis et al., 2019; Salas et al., 2018; Tolstenkov et al., 2025; Yang et al., 2026). Upon adherence to an appropriate substrate, metamorphosis can be triggered through mechanosensory cues (Hozumi et al., 2025; Totsuka and Hotta, 2025; Wakai et al., 2021). Settlement is further enhanced or suppressed by chemical cues sensed through multimodal sensory neurons located in or near the anterior adhesive papillae of the larva (Hoyer et al., 2024; Yang et al., 2026).

Although tunicates have been studied as model organisms for cell, developmental, and organismal biology for over 100 years (Conklin, 1905; Lemaire, 2009), various molecular techniques have been more recently adapted to studying the *Ciona* CNS (Olivo et al., 2021). Of note, CRISPR/Cas9 is now routinely used to study gene function in *Ciona* and other tunicates (Pennati et al., 2024). Here we publish the design and validation of a handful of novel single-chain guide RNAs (sgRNAs) for tissue-specific CRISPR/Cas9-mediated mutagenesis in the larval CNS of *Ciona robusta*. Their targets comprise a variety of conserved transcriptional regulators of neurodevelopment, in addition to two conserved effectors of sensory/neural cell function. Most of these genes have not been previously knocked out in *Ciona* using CRISPR/Cas9. These reagents are being made freely available to the *Ciona* community, in the hopes that they can be readily used to further study the larval nervous system.

## Methods and results

We sought to design and validate sgRNAs targeting several *C. robusta* genes involved in neural development and function. To start, we selected 6 transcription factor genes with well-documented expression patterns in *Ciona* development (Imai et al., 2004; Imai et al., 2006; Imai et al., 2009), encoding a mix of early-expressed regulators (*Cdx, Foxb, Sox1/2/3)* and later-onset ones (*Dmbx, Engrailed, Mnx)*. Additionally, we designed sgRNAs targeting the *Tyrosinase* (*Tyr*) gene, which encodes the rate-limiting enzyme for melanin biosynthesis in pigmented cells associated with photo- and geo-sensory organs of the larva. *Tyr-*targeting sgRNAs have already been designed (Pickett and Zeller, 2018; Song et al., 2022; Vitrinel et al., 2023), though these have not been formally validated by target amplicon next-generation sequencing, to our knowledge. Finally, we also designed sgRNAs targeting *Slc18a3* (commonly referred to as *Vesicular acetylcholine transporter*, or *VAChT*). This gene part of a highly conserved “cholinergic locus” that also encodes for Choline acetyltransferase (ChAT), with both *VAChT* and *ChAT* transcripts sharing a first exon (Kratsios et al., 2012). In *Ciona, VAChT/ChAT* is expressed in motor neurons and other cholinergic neurons of the CNS (Horie et al., 2010; Imai et al., 2009; Jokura et al., 2020; Nishino et al., 2011; Takamura et al., 2010; Yoshida et al., 2004).

Candidate sgRNAs were identified using the online webtool CRISPOR (Concordet and Haeussler, 2018), according to our recent guidelines (Popsuj et al., 2024). Where possible, sgRNAs with predicted Doench ‘16 efficacy scores >50 were selected. For *Dmbx*, suitable sgRNAs were limited due to its very small exons and a paucity of highly scored sgRNA candidates. The sgRNA expression plasmids (containing the small U6 RNA promoter) were constructed using either older oligonucleotide- or PCR-based subcloning protocols (Stolfi et al., 2014) or a newer approach based on *de novo* gene synthesis and custom cloning services (Twist Bioscience)(Johnson et al., 2023). In all, 25 sgRNAs targeting the 8 genes were tested by a standard, commercially available Illumina-based, next-generation target site amplicon sequencing service (Amplicon-EZ by Genewiz) as previously described (Popsuj et al., 2024). The protocol for sgRNA validations can be found at: http://dx.doi.org/10.17504/protocols.io.14egnr97yl5d/v1

Amplicon sequencing revealed indels at the target site for all sgRNAs tested, with varying indel frequency, or mutation efficacy. Those sgRNAs with the best, or acceptable measured indel rates are shown according to their target location in each locus, along with their corresponding amplicon sequencing indel plots (**Figures 1-3**). These are the sgRNAs we would recommend using to disrupt these genes. All genes had at least one sgRNA with efficacy >30%, again with the exception of *Dmbx*. Although we designed and tested 5 sgRNAs against this gene, the highest efficacy achieved was 25% with one particular sgRNA (**Figure 2**). All sgRNA sequences, PCR primer sequences, and the predicted and measured efficacies of each sgRNA are presented here (**Figure 4, Supplemental Table 1, Supplemental Sequences File**). Some of the measured efficacies were approximated based on the resulting indel plot, though naturally occurring indels and SNPs prevented the automated return of a precise efficacy value.

**Figure 1.**
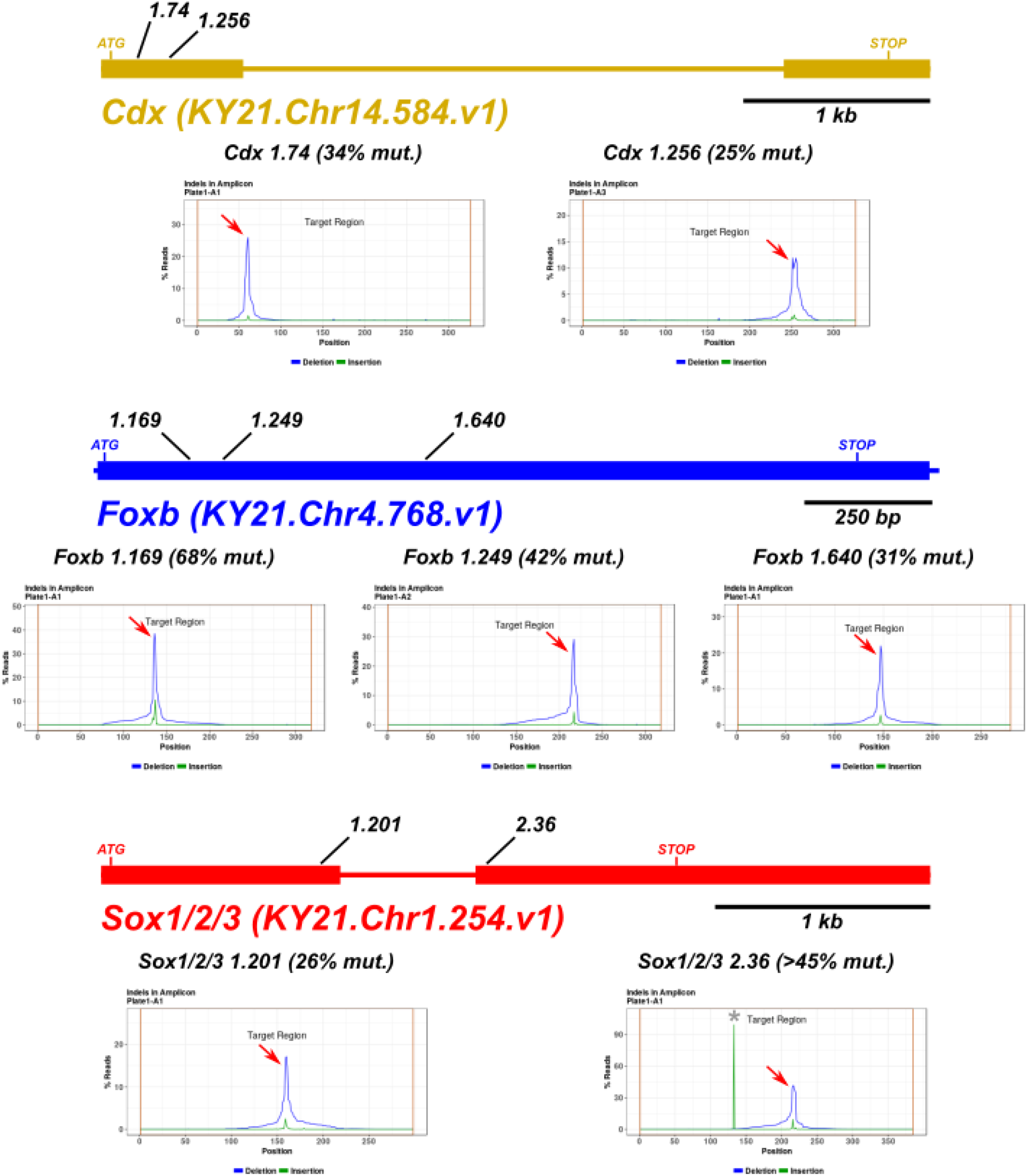
*Cdx, Foxb*, and *Sox1/2/3* loci and sgRNA validation results. Selected sgRNA target sites are indicated, along with Amplicon-EZ (Genewiz) indel plots and estimated mutagenesis rates. Red arrows indicate CRISPR-generated indel peaks. Grey asterisks indicate naturally-occurring indels. Mutagenesis rates are approximated for plots in which naturally occurring indels confounded automatic rate estimation.

**Figure 2.**
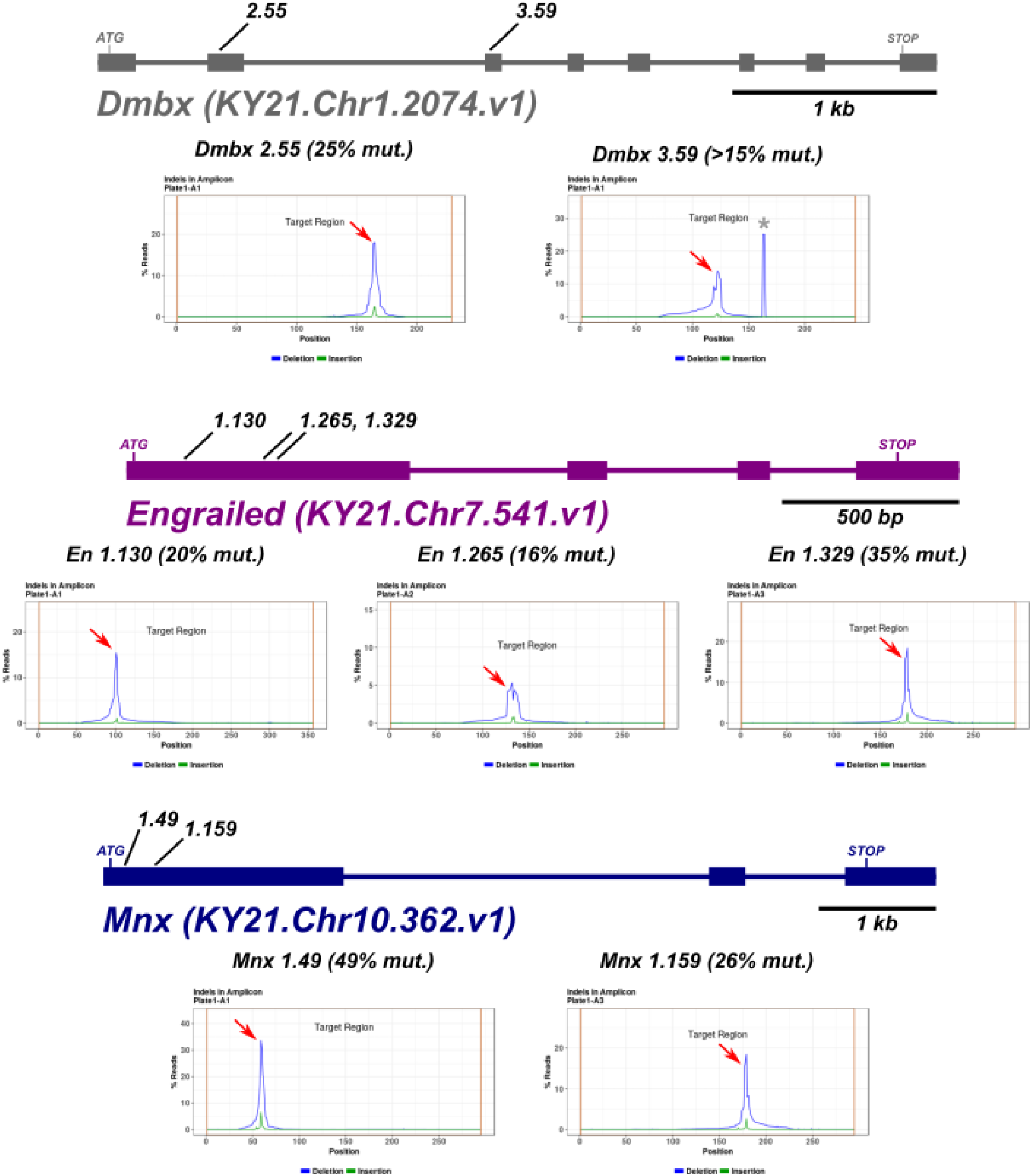
*Dmbx, Engrailed*, and *Mnx* loci and sgRNA validation results. Selected sgRNA target sites are indicated, along with Amplicon-EZ (Genewiz) indel plots and estimated mutagenesis rates. Red arrows indicate CRISPR-generated indel peaks. Grey asterisks indicate naturally-occurring indels. Mutagenesis rates are approximated for plots in which naturally occurring indels confounded automatic rate estimation.

**Figure 3.**
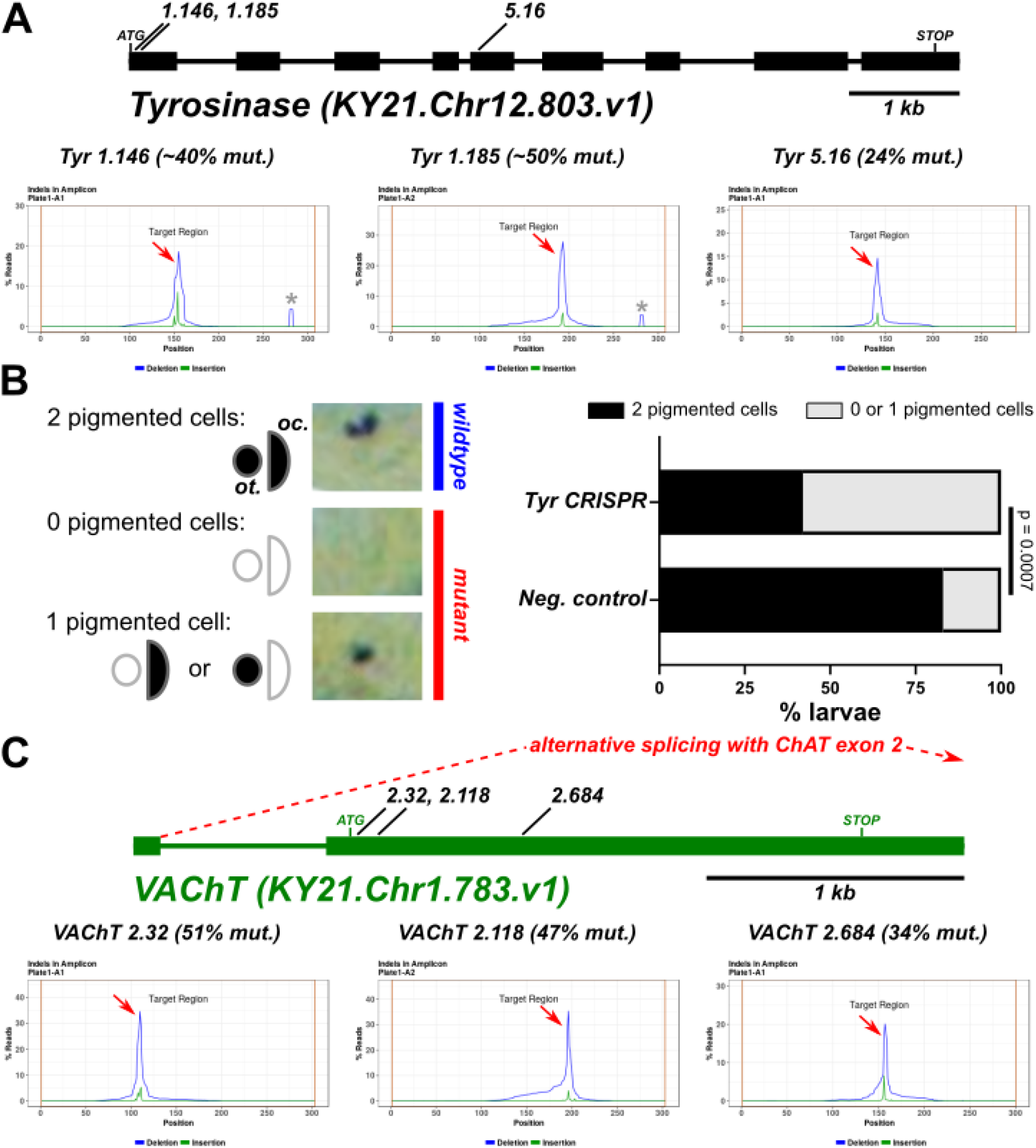
*Tyrosinase* and *VAChT* loci and sgRNA validation results. A) Selected *Tyrosinase* sgRNA target sites are indicated, along with Amplicon-EZ (Genewiz) indel plots and estimated mutagenesis rates. Mutagenesis rates are approximated for plots in which naturally occurring indels confounded automatic rate estimation. B) Left: diagram of simple pigmentation assay (see text for details). Right: results of pigmentation assay of *Tyr CRISPR* larvae using sgRNAs 1.185 and 5.16 together (40 µg each) in combination with 50 µg *Sox1/2/3>Cas9::Geminin*^*Nterminus*^. (Pennati et al., 2024; Stolfi et al., 2014). CRISPR larvae (n = 45) were compared to negative control larvae (n = 29) electroporated with 80 µg of the “control” sgRNA plasmid instead (Stolfi et al., 2014). Plasmid DNA amounts indicated per 700 µl of total electroporation volume. Statistical significance determined using Fisher’s exact test. C) Selected *VAChT* sgRNA target sites indicated, along with indel plots. Dashed red line indicates alternative splicing of first exon with remaining exons encoding choline acetyltransferase (ChAT) out of view, as part of a conserved cholinergic locus. Red arrows indicate CRISPR-generated indel peaks. Grey asterisks indicate naturally-occurring indels.

**Figure 4.**
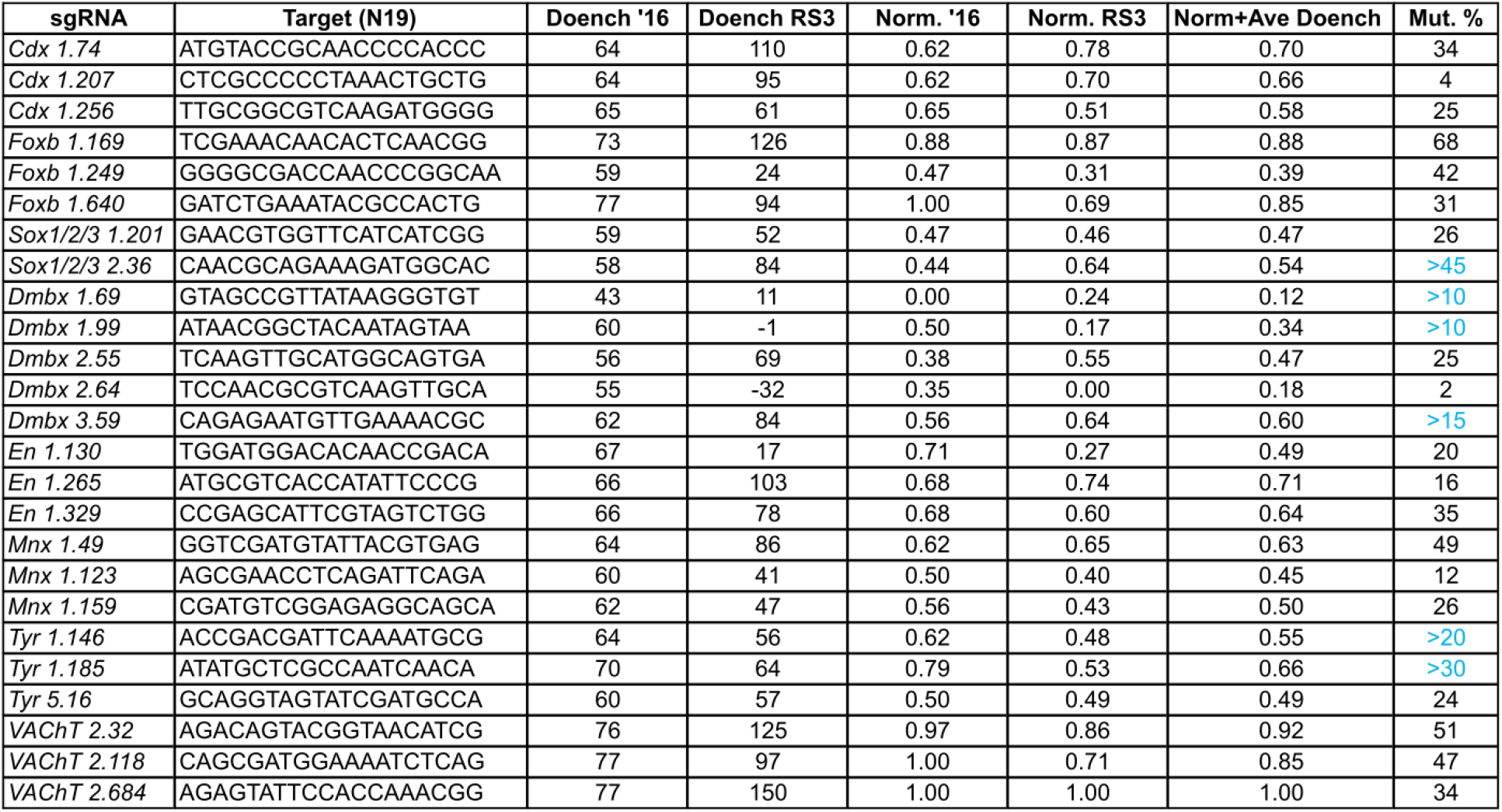
Table of all sgRNAs designed and tested in this study. All sgRNA target sequences tested in this study, along with predicted efficacy scores from CRISPOR, namely Doench ‘16 and Doench Ruleset 3 (RS3). Norm. ‘16 and Norm. RS3 represents normalized Doench ‘16 and Doench RS3 scores, in which the represented ranges (not absolute theoretical score ranges) are converted to a scale from 0 to 1. Norm+Ave = normalized Doench ‘16 and RS3 scores averaged. Mut. % = mutagenesis efficacy as measured by Amplicon-EZ sequencing. Mut. % scores in blue represent efficacies estimated from size of indel peak, and not the automated efficacies as given by automated Amplicon-EZ analysis (black Mut. % scores).

To further validate the efficacy of *Tyr* sgRNAs, we performed a simple cell pigmentation assay (**Figure 3B**). We combined sgRNA expression vectors 1.185 and 5.16, which target separate exons, and co-electorporated them with *Sox1/2/3>Cas9::Geminin*^*Nterminus*^ to knock out *Tyr* in the ectoderm from which the pigment cells are derived. While wild-type larvae typically have two pigment cells (1 ocellus pigment cell and 1 otolith), loss of Tyrosinase function results in loss of pigmentation (Jiang et al., 2005; Pickett and Zeller, 2018). Indeed, we found a statistically significant increase in the frequency of larvae with zero or only one pigment cell. The latter is expected in cases of left/right mosaic knockouts. The ∼50% reduction in pigment cell frequency was around that expected given the sgRNA efficacies we measured by amplicon sequencing (**Figure 3A**).

CRISPOR recently introduced the use of an updated Doench ‘16 algorithm, named “Doench Ruleset 3” (RS3)(DeWeirdt et al., 2022). Whereas we typically avoid Doench ‘16 scores <50, RS3 scores span a larger range of values. However, our initial sgRNA designs did not take Doench RS3 into account. We therefore sought to re-evaluate the correlation between these predictive algorithms and the actual mutagenesis efficacies. In our small dataset, there was moderate correlation between predicted scores and measured mutagenesis efficacies, with slightly higher Pearson correlation coefficients specifically for the RS3 algorithm (**Figure 5**). The correlation was slightly higher when comparing measured efficacies to the average of both Doench scores after normalization (“Doench Norm+Ave”). We therefore recommend that both Doench scores (‘16 and RS3) be taken into consideration when selecting candidate sgRNAs for CRISPR in *C. robusta*, though RS3 might be the more reliable predictive algorithm going forward. Future studies using an unbiased set of predicted sgRNAs will be needed to more thoroughly compare these algorithms, bearing in mind that the sgRNAs analyzed here were already selected for high Doench ‘16 scores.

**Figure 5.**
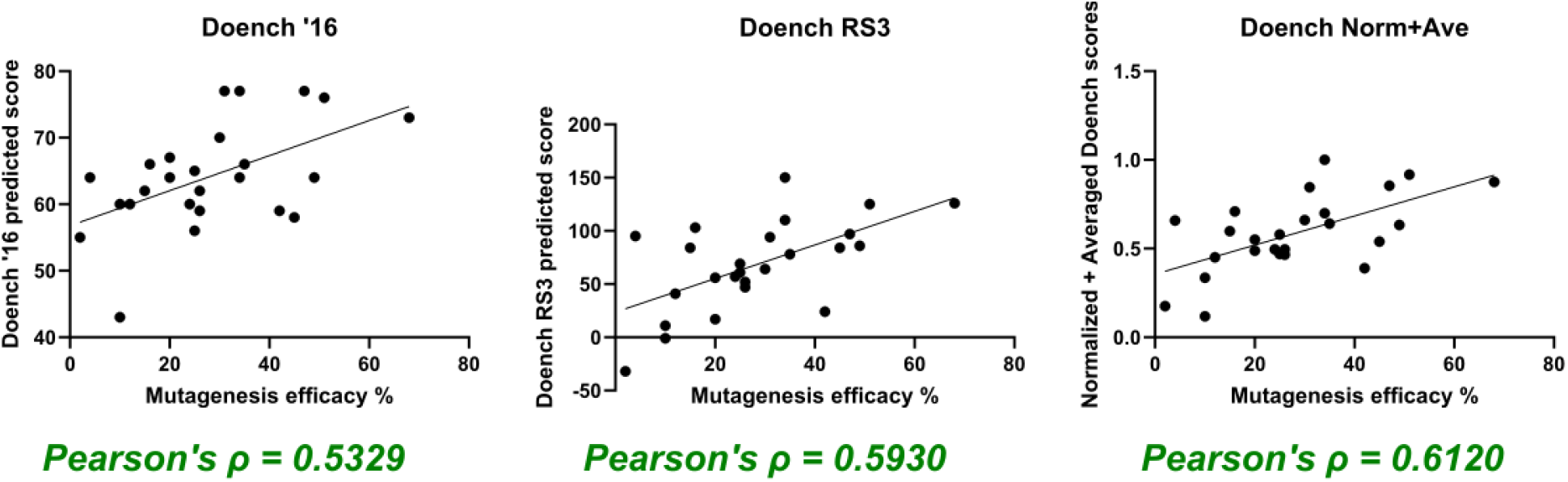
Plotting predicted and measured mutagenesis efficacies. Plots comparing predicted efficacy scores (Doench ‘16, Doench Ruleset 3/RS3, or normalized/averaged Doench scores) to actual mutagenesis efficacies as measured/estimated by Amplicon-EZ sequencing. All data according to the table in Figure 4. Pearson’s correlations calculated and shown for each set of predictions.

## Conclusion

We have designed and tested novel sgRNA expression plasmids targeting 8 different neurodevelopmental genes in the model tunicate *Ciona robusta*. We are publishing and sharing these immediately, in the hopes that they will be of use for the larger *Ciona* research community. These and other published plasmids (e.g. Cas9 expression plasmids) are available upon request. We are also happy to share access to our onboarded sgRNA and Cas9 expression at Twist Bioscience, for easy custom synthesis and cloning of new sgRNAs and Cas9 vectors. Finally, comparing predicted and measured indel efficacy rates suggests that the new “Doench Ruleset 3” score in the online prediction tool CRISPOR might be the preferred algorithm to consider when picking new sgRNAs for CRISPR in *Ciona*.

## Supporting information

Supplemental Sequences

## Acknowledgments

This work was funded by NIH grants R35GM158421 and R01HD104825 to A.S. We thank Dexter Dean and past and current members of the lab for helpful feedback and technical assistance.

