## Supplemental Sequences for "Validated CRISPR/Cas9 guide RNAs targeting neurodevelopmental genes in the tunicate *Ciona robusta*"

**Supplemental Sequences File**

**New single-chain guide RNAs (sgRNAs) validated in this study**

Cdx 1.74

**GATGTACCGCAACCCCACCC (G+N19)**

Cdx 1.207

**GCTCGCCCCCTAAACTGCTG (G+N19)**

Cdx 1.256

**GTTGCGGCGTCAAGATGGGG (G+N19)**

Dmbx 1.69

**GGTAGCCGTTATAAGGGTGT (G+N19)**

Dmbx 1.99

**GATAACGGCTACAATAGTAA (G+N19)**

Dmbx 2.55

**GTCAAGTTGCATGGCAGTGA (G+N19)**

Dmbx 2.64

**GTCCAACGCGTCAAGTTGCA (G+N19)**

Dmbx 3.59

**GCAGAGAATGTTGAAAACGC (G+N19)**

En 1.130

**GTGGATGGACACAACCGACA (G+N19)**

En 1.265

**GATGCGTCACCATATTCCCG** **(G+N19)**

En 1.329

**GCCGAGCATTCGTAGTCTGG (G+N19)**

Foxb 1.169

**GTCGAAACAACACTCAACGG (G+N19)**

Foxb 1.249

**GGGGGCGACCAACCCGGCAA** **(G+N19)**

Foxb 1.640

**GGATCTGAAATACGCCACTG (G+N19)**

Mnx 1.49

**GGGTCGATGTATTACGTGAG (G+N19)**

Mnx 1.123

**GAGCGAACCTCAGATTCAGA (G+N19)**

Mnx 1.159

**GCGATGTCGGAGAGGCAGCA (G+N19)**

Sox1/2/3 1.201

**GGAACGTGGTTCATCATCGG (G+N19)**

Sox1/2/3 2.36

**GCAACGCAGAAAGATGGCAC (G+N19)**

Tyr 1.146

**GACCGACGATTCAAAATGCG (G+N19)**

Tyr 1.185

**GATATGCTCGCCAATCAACA (G+N19)**

Tyr 5.16

**GGCAGGTAGTATCGATGCCA (G+N19)**

VAChT 2.32

**GAGACAGTACGGTAACATCG (G+N19)**

VAChT 2.118

**GCAGCGATGGAAAATCTCAG (G+N19)**

VAChT 2.684

**GAGAGTATTCCACCAAACGG (G+N19)**

**Primers for NGS validation of sgRNAs**

| **Amplicon** | **Forward primer** | **Reverse primer** |
| --- | --- | --- |
| Cdx exon 1 | TGAGAAACGCACAATGTTATG | AGTTCGAATTCATGTCCCATG |
| Dmbx exon 1 | TTCGTGCAATGTCAGTGTTC | GCACGCCTTTGAGTTACAAT |
| Dmbx exon 2 | ATTGTAACTCAAAGGCGTGC | AGATTGCCAAACTTTCTCGC |
| Dmbx exon 3 | TTGTGCATTGCTTCCGAAAC | GGTGAAACACGGTCGTTTTG |
| Engrailed exon 1 | CGAAATACAAGGTTGCCCAAC | GGATCACGCACATCGTTTATC |
| Foxb 1.169, 1.249 | ACCAGGACGAGACTCATATG | CATTTTGAATCACAGACCGTC |
| Foxb 1.640 | GCCATAACTTATACCAGGCAC | GGTTAGGTGCGTTTGCATTG |
| Mnx exon 1 | CTTCTATCGCATTGAACCTTTGA | TGAAAACTTGGCTCGTTTCG |
| Sox1/2/3 exon 1 | ACTGGATGCAGATCCCATAC | TAAGTGGCGTATGCTAACAC |
| Sox1/2/3 exon 2 | CGAACAATGCGATTCTAGACAC | CTCTTTCTTAAGGATCGCCTTC |
| Tyr exon 1 | GGCACAAGTTATGTCTTGTATT | CTATTATGCCTATATTCCGCGAG |
| Tyr exon 5 | GAAAACAGAGCTGAACTCGTC | TGGCTGATAAGTGTTACGTTATC |
| VAChT 2.32, 2.118 | GCCTTTTGCAGATTACATCTG | GAAAGCACTGCGGACTTTG |
| VAChT 2.684 | CAGAGGGCTTTGCTGTATTG | GCTTATAGATCGGTGTACCAAC |

**ID numbers of gene sequences targeted in this study**

| **Gene name** | **KH ID** | **KY21 ID** | **ANISEED ID** |
| --- | --- | --- | --- |
| Cdx | C14.408 | Chr14.584 | Cirobu.g00003840 |
| Dmbx | C1.1212 | Chr1.2074 | Cirobu.g00000237 |
| Engrailed (En) | C7.431 | Chr7.541 | Cirobu.g00008277 |
| Foxb | C4.341 | Chr4.768 | Cirobu.g00006425 |
| Mnx | L128.12 | Chr10.362 | Cirobu.g00010957 |
| Sox1/2/3 | C1.99 | Chr1.254 | Cirobu.g00001283 |
| Tyrosinase (Tyr) | C12.469 | Chr12.803 | Cirobu.g00003025 |
| VAChT | C1.498 (v1) | Chr1.783 (v1) | Cirobu.g00000742 |
